# Discovery of a new but established population of the guppy in Germany

**DOI:** 10.1101/2022.02.01.478389

**Authors:** David Bierbach, Christopher Schutz, Nils Weimar, Alessandra Escurra Alegre, Fritz A. Francisco, Serafina Wersing, Olivia L. O’Connor, Michael Kempkes, Udo Rose, Friedrich Wilhelm Miesen, Marie Habedank, Jonas Jourdan, Jana Kabus, Delia Hof, Simon Hornung, Sebastian Emde, Leon Rüffert, Juliane Lukas

## Abstract

Feral populations of tropical fish species in temperate climates like Central Europe are a rare but repeatedly observed phenomenon. Due to the influence of industrial or geothermal heated water, released tropical fish may be able to survive harsh winter conditions. Here we characterize a newly discovered thermally polluted river, with an established population of the guppy (*Poecilia reticulata*) co-occurring with native species. Through a mark-recapture approach, we estimated the population size of the guppies close to the warm water inflow to be around 2000 individuals during summer and we further provide descriptive demographics of this population which allow us to assume it is well established in that river. Further, we found some of the sampled specimen being parasitized by *Camallanus* roundworms, thus showing the guppies’ host potential for this genus of internal parasites. The popularity and widespread distribution of guppies as ornamental fish often leads to their intentional or unintentional release into the wild where they are often pioneer species in anthropogenically heavily modified habitats. Guppies threaten native species through niche competition and transmission of diseases. Accordingly, early awareness and knowledge on the status of non-native populations is crucial for effective management strategies.

## Introduction

Global warming and biological invasions are two of the most pressing extinction threats in aquatic ecosystems (Sala et al. 2000). Our understanding of their complex interplay and, potentially, detrimental synergies between these two extinction drivers is still limited - despite urgent need for improved management approaches and enhanced conservation efforts to mitigate, prevent, or, ultimately, reverse global freshwater biodiversity loss (Keller et al. 2011, Sorte et al. 2013, Latombe et al. 2017). In order to investigate these effects, thermally altered freshwaters have been proposed as promising semi-natural research laboratories to address the underlying challenges and questions (Lukas et al. 2021). Water bodies in the temperate zones that receive natural and/or anthropogenic warm water influxes represent unique temperature refuges that allow non-native species (especially those of tropical origin) to establish self-sustaining populations (Langford 1990). Besides their suitability for semi-natural experiments on migration, spread dynamics and adaptability of non-native species within observable temporal and spatial scales (O’Gorman et al. 2014, Lukas et al. 2021), these populations are further known to introduce non-native diseases (Emde et al. 2015). Knowledge of the presence of self-sustaining tropical fish populations in temperate areas is thus not only representing research opportunities but is also mandatory for management actions to be taken (Latombe et al. 2017). In Germany, there are 3 recent sites known that harbor stable, self-sustaining populations of tropical fish species (Jourdan et al. 2014, Lukas et al. 2017a, Lukas et al. 2017b, Kempkes et al. 2018, Lukas et al. 2021). Here we report on a newly discovered population of feral guppies close to but independent of the well-reported Gillbach population in western Germany (Jourdan et al. 2014).

## Results

Our sampling took place at the river ‘Kleine Erft” located within the authority of the City of Bergheim (50° 56’ 31.87” N, 6° 39’ 34.82”; federal state of North Rhine-Westphalia, Germany, Figure 1B). Casual sightings of guppies by one of the authors (U.R.) led to the first intensive sampling in this area in July 2021 by all of the authors and a subsequent sampling in September 2021. The “Kleine Erft” river receives warm water influx from a nearby industrial area through a pipe of ca. 1 meter diameter. We measured the river’s physico-chemical water parameters 5 m upstream and 160 m downstream of the warm water influx (pH, EC, water temperature) according to the manufacturers’ protocols and recommendations (Manufacturer: HANNA). Water temperatures 5 m upstream the inflow pipe was 18.8°C while the warm water inflow increased the temperatures immediately to 25.8°C. The temperatures stayed above the colder upstream values for more than 150 m (Figure 1B, see Table S1). pH and EC were relatively constant along the investigated river stretch: pH-range: 7.9-8.0, EC range: 1.05 - 1.12 mS (see details at supplemental Table S1). For a long-term temperature profile we placed HOBO pendant data loggers (Onset Computer Corp.) in close vicinity of the warm water influx from July to September 2021. For a comparison, we also present data from loggers placed in the Gillbach that is ca. 10 km away and which is known for its warm water regime and guppy occurrences (Jourdan et al. 2014). Our long-term temperature recordings found the river ‘Kleine Erft’ to have on average 5.3°C ± 2.2°C lower daily mean temperatures compared to the Gillbach (Figure 1A). This is apparently due to the mixing of warm water from the influx and colder stream water in the Kleine Erft compared to a pure warm water source in the Gillbach. In both rivers, water temperature was affected by weather but also random, most likely discharge-related variations (Figure 1A).

**Figure 1.**
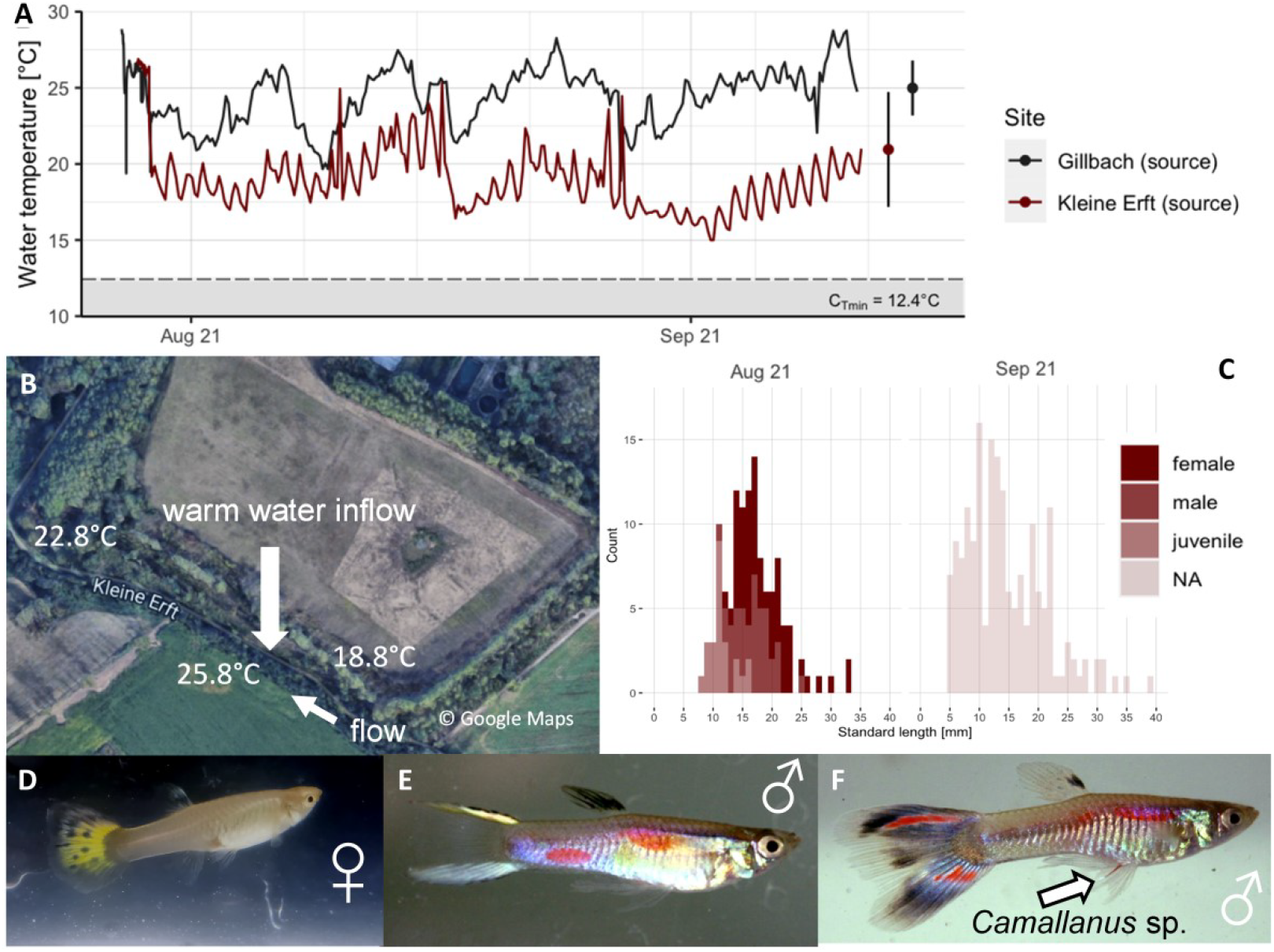
(A) Long-term measures at the warm water influx for the “Kleine Erft” as well as the Gillbach in 2021 (B) Location of the river “Kleine Erft” and the warm water inflow pipe (arrow). Here the manually measured temperatures during the July sampling are indicated. (C) Body size distribution of the found guppies (*Poecilia reticulata*) during both sampling events. Please note that sexes were only distinguished in July. (D) Female with ornamental-bred color-pattern in the caudal fin. (E & F) Examples of male color morphs found at the Kleine Erft. Male shown in (F) with *Camallanus* roundworm (arrow).

Fish and macroinvertebrates were sampled using dip-nets (mesh size 2 mm), 2-person seines (mesh size 4 mm) as well as electro-fishing gears (EFGI 650, Bretschneider). We spent 4 hours sampling at the site in July (all gears) and another 4 hours in September (only seine and dip-net fishing), sampling a stretch 200 m downstream and 50 m upstream the warm water inflow pipe. We found 9 fish species in the ‘Kleine Erft’, 2 of which are considered non-native with the guppy being the only species from the tropics (Table 1). European chub was the dominant native species with 67 specimens caught. Further, non-native shrimps (*Neocardinia davidii*) and water snails (*Melanoides tuberculata*) were found in high densities. Please note that guppies and similarly small specimen were not caught with electro-fishing gears thus we did not use this gear for the recapture trip in September (see below).

**Table 1:** Results from fish stock survey at river Kleine Erft. Shown are found species with abundancies and body sizes as well as sampling gears. Please note that guppy body sizes are measured as SL (standard length) due to the often elongated male caudal fins (see Fig. 1E&F).

| Species | common name | Native/non-native | Average TL [cm] | Frequency [ind] | Min TL [cm] | Max TL [cm] | Sampling method |
| --- | --- | --- | --- | --- | --- | --- | --- |
| *Alburnus alburnus* | Common bleak | Native | 2.8 | 15 | 1.2 | 15 | E-Fishing / seine |
| *Barbatula barbatula* | Stone loach | Native | 10 | 1 | 10 | 10 | E-Fishing |
| *Cyprinus carpio* | Common carp | Non-native | - | 1 | - | - | Sighted |
| *Gasterosteus aculeatus* | Three-spined stickleback | Native | 2 | 3 | 1.7 | 2.5 | Seine |
| *Gobio gobio* | Gudgeon | Native | 3.5 | 18 | 1.6 | 6.4 | E-Fishing / seine |
| *Perca fluviatilis* | European perch | Native | 30 | 1 | 30 | 30 | E-Fishing |
| *Poecilia reticulata* | Guppy | Non-native | SL:1.5 | 294 | SL:0.5 | SL:3.9 | Seine |
| *Rutilus rutilus* | Common roach | Native | 20.8 | 6 | 15 | 25 | E-Fishing |
| *Squalius cephalus* | European chub | Native | 21 | 67 | 10 | 40 | E-Fishing |

Guppies showed phenotypes containing many ornamental bred features like female coloration (Figure 1D) and males with elongated and pigmented caudal fins (figure 1 E&F), suggesting that the founder animals of this population were ornamental fish released by humans.

From the caught guppies, we took 40 adult individuals to our laboratory at HU Berlin for further analysis during the July sampling and 25 different specimen were color-tagged using VIE (Jourdan et al. 2014) and afterwards released on site. All guppies were measured for standard length using either pictures taken on millimeter paper or rulers on site. During the second sampling in September, we used dip-nets and seines to recapture marked guppies for population size estimates using the R package ‘Rcapture’ (Baillargeon and Rivest 2007) assuming a closed population (Mt model, see supplemental Table S2).

From the initially marked fish (15 males and 10 females), we recaptured 2 specimen in September (1 male and 1 female) in a catch of 176 guppies during the second sampling. When we assume no deaths or migration having occurred between samplings, this leads to an overall population size estimate of 2376 (± 1607) guppies in the river Kleine Erft.

Body sizes of found guppies comprised both juveniles and adults showing the self-sustaining character of this population (Figure 1C). For guppies sampled in July, we established the sex of caught adults as far as possible (SL above 10 mm, Gonopodium visible in males) which revealed a female biased sex-ratio of 57f:39m (1 to 0.68). Interestingly, two of the males that were taken to HU Berlin alive were parasitized by a *Camallanus* roundworm (figure 1 F, region of Gonopodium, indicated with arrow). This is particularly important as it shows that guppies of the Kleine Erft population could serve as a host for native European *Camallanus* species as well as non-native *C. cotti*. Full methods are provided as supplemental information.

## Discussion

Here we report on the discovery of a yet not reported feral population of the guppy (*P. reticulata*) inhabiting a thermally altered river in western Germany. Water temperatures around the warm water influx were increased for a stretch of at least 150 m from 18.8°C to 25.8°C, rendering this river a suitable habitat for the tropical guppy as the lower critical minimum temperature for feral guppies in Germany was reported to be 12.4°C (Jourdan et al. 2014). While the guppy is the only tropical fish found so far in the ‘Kleine Erft’ river, body sizes of found specimen as well as population size estimates point towards a self-sustaining, thus established population. Please note that the mark-recapture-based population estimates are very rough due to the low numbers of marked and recaptured individuals. In contrast to closely related species such as *Gambusia holbrooki* which is already established throughout Southern Europe (Jourdan et al. 2021), the guppy is not expected to spread outside of warm-water inputs in Central Europe. However, the co-occurrence of native European fish species along with the records of *Camallanus* parasites in several guppies recommends a continuous monitoring of this population (Latombe et al. 2017) as non-native parasites using guppies as hosts have been shown to switch also to native fish species in this area of Germany posing unpredictable threats to native European fish stocks (Emde et al. 2015).

## Supporting information

Full methods and additional results

Data on fish sampling

## Acknowledgement

We thank the Erftverband as well as the LANUV for their support and the provisioning of fishing permits. This research was financially supported by overhead funds from the Deutsche Forschungsgemeinschaft (DFG) as well as the German Ichthyological Society (GfI).

## Author contributions

All authors contributed to the sampling in July; CS and NW performed sampling in September; JL analyzed the data; DB wrote the manuscript with input from all authors.

