## Supplementary material for "Discovery of a new but established population of the guppy in Germany": Full methods and additional results

**Sampling area and water parameters**

Our sampling took place at the river ‘Kleine Erft” located within the authority of the City of Bergheim (50° 56' 31.87" N, 6° 39' 34.82"; federal state of North Rhine-Westphalia, Germany, Figure 1B). Casual sightings of guppies by one of the authors (U.R.) led to the first intensive sampling in this area in July 2021 by all of the authors and a subsequent sampling in September 2021. The “Kleine Erft” river receives warm water influx from a nearby industrial area through a pipe of ca. 1 meter diameter. We measured the river’s physico-chemical water parameters 5 m upstream and 160 m downstream of the warm water influx (pH, EC, water temperature) according to the manufacturers’ protocols and recommendations (Manufacturer: HANNA). For a long-term temperature profile we placed HOBO pendant data loggers (Onset Computer Corp.) in close vicinity of the warm water influx from July to September 2021. For a comparison, we also present data from loggers placed in the Gillbach that is ca. 10 km away and which is known for its warm water regime.

Table S1: Water temperature, pH and EC measured at “Kleine Erft” in Juli 2021.

| **Distance from source [m]** | **River side** | **Water temperature [C°]** | **pH** | **EC [mS]** |
| --- | --- | --- | --- | --- |
| 0 | Warm water influx | 25.8 | 7.9 | 1.18 |
| 3 | right | 24.7 | 7.9 | 1.15 |
| 3 | left | 24.0 | 7.9 | 1.05 |
| 8 | right | 22.4 | 8.0 | 1.07 |
| 8 | left | 23.8 | 8.0 | 1.12 |
| 11 | right | 23.2 | 8.0 | 1.07 |
| 11 | left | 24.0 | 7.9 | 1.12 |
| 15 | right | 22.7 | 8.0 | 1.10 |
| 15 | left | 23.8 | 8.0 | 1.12 |
| 18 | right | 23.2 | 8.0 | 1.11 |
| 18 | left | 23.9 | 8.0 | 1.12 |
| 22 | right | 23.6 | 8.0 | 1.11 |
| 22 | left | 23.5 | 8.0 | 1.11 |
| 25 | right | 23.5 | 8.0 | 1.10 |
| 25 | left | 22.8 | 8.0 | 1.09 |
| 150 | right | 22.8 | 8.0 | 1.08 |
| 150 | left | 22.8 | 8.0 | 1.07 |

**Fish stock survey**

Fish and macroinvertebrates were sampled using dip-nets (mesh size 2 mm), 2-person seines (mesh size 4 mm) as well as electro-fishing gears (EFGI 650, Bretschneider). We spent 4 hours sampling at the site in July (all gears) and another 4 hours in September (only seine and dip-net fishing), sampling a stretch 200 m downstream and 50 m upstream the warm water inflow pipe. Fish species were determined to species level using Kottelat and Freyhof, J. (2007).

**Mark-Recapture population size estimation of guppy population**

From the caught guppies, we took 40 adult individuals to our laboratory at HU Berlin for further analysis during the July sampling and 25 different specimen were color-tagged using VIE (Jourdan et al. 2014) and afterwards released on site. All guppies were measured for standard length using either pictures taken on millimeter paper or rulers on site. During the second sampling in September, we used dip-nets and seines to recapture marked guppies for population size estimates using the R package ‘Rcapture’ (Baillargeon and Rivest 2007) assuming a closed population (Mt model, see supplemental Table S2).

Table S2: Results from ‘Mark-Recapture’ models using the R package ‘Rcapture’ (Baillargeon and Rivest 2007)

| **Model** | **abundance** | **SEM** | **df** | **AIC** | **BIC** |
| --- | --- | --- | --- | --- | --- |
| M0 | 5151.1 | 3570.6 | 1 | 144.107 | 150.714 |
| Mt | 2376.0 | 1607.5 | 0 | 20.675 | 30.585 |
